# On the synchronized failure of global crop production

**DOI:** 10.1101/287391

**Authors:** Zia Mehrabi, Navin Ramankutty

## Abstract

Multiple breadbasket failure is a risk to global food security. However, there are no global analyses that have assessed if global food production has actually tended towards synchronized failure historically. We show that synchronization in production for major commodities such as maize and soy has declined in recent decades, but that increased synchrony, when present, has had marked destabilizing effects. Under the hypothetical case of a synchronized failure event, we estimate simultaneous global production losses for rice, wheat, soy and maize between −18% and −36%. Our results show that maintaining asynchrony in the food system and mitigating instability through food storage in good years, both require a central place in discussions of future food demand under mean climate change, population growth and consumption trends.

## Introduction

Over the past century humanity has experienced considerable climatic, economic and political shocks to the food system (1–8). These shocks have been associated with regional food shortages (5, 9, 10), price spikes (3, 11) and food insecurity (1, 4, 12, 13). In recent years, scientists, governments and the insurance industry have joined forces in an attempt to identify the future risk posed by food system shocks (4, 14). A key concern is that if co-occurring shocks were to hit multiple breadbaskets in the future, this would lead to large losses in food production, and, in some cases, to civil unrest (4, 14, 15). Notwithstanding the interest in this area, and recent work to identify the global impacts of isolated extreme weather disasters on crop production (2), and conflict (15–17), it is currently unknown if the food system has actually tended towards synchronized failure in recent history. Critically, a better understanding of the historical stability of food production might help to better anticipate the expected losses under synchronized failure in the future, and devise strategies to mitigate potential losses.

Here we present an analysis of the stability of global production for four major commodities (maize, rice, soy and wheat, making up ~60% of global production) over 1961-2008. We identify which locations on the planet have historically reduced or increased the inter-annual variation in production at the global level, and perform diagnostics to assess if the food system has shown signs of increasing synchrony or instability in production in recent decades. We then use the empirical variation in historical production trends, which contain information on the impact of many different production shocks, including, but not limited to, natural disasters and systemic economic breakdowns, to estimate the maximum observed inter-annual deficits in global crop production, and the expected inflation of these global deficits under synchronized production failure. Finally, we explore the potential impact of four radical mitigation strategies – closing production gaps, closing production ceilings, global adoption of more resilient cropping systems, and focused efforts to adopting resilient cropping systems in the world’s major breadbaskets - on offsetting the expected losses under the empirically grounded worst case scenario of synchronized failure.

## Results

### Mapping local contributions to global variance in production

Local contributions to the inter-annual variance in global crop production 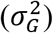 over 1961-2008 are shown in Figure 1. We removed the temporal trends in production from the data, so that only the year-to-year variation is represented: gains and losses that would otherwise be swamped by the net changes in production due to technology improvement over this time period (18, 19). Not surprisingly, the major breadbaskets contain locations that have historically made annual crop production more variable (i.e. increase 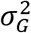). For example, we see locations in the mid-west US increasing the variance in global trends in maize, with the worst 100km x 100km area production grid cells accounting for 4.1% of the variation in global production trends over the period of 1961-2008; and for soy by up to 3.1%. We also see individual locations in Northern India accounting for as much as 2.6% of the inter-annual variation in global rice trends. We also identify that spatial compensation has occurred over the same period, albeit with smaller effects and more restricted extents. For example, locations in South America and India have played important roles in reducing variation in global soy trends by as much as 0.4%. Importantly, whilst locations in Eastern Europe do show evidence of increasing the variation in wheat trends by up to ~1.3%, as might be expected from the importance of this breadbasket, our analysis clearly shows that the global variation of wheat production depends much less on specific locations, countries or regions, than any of the other three major crop commodities (Figs 1 A-D).

**Figure 1.**
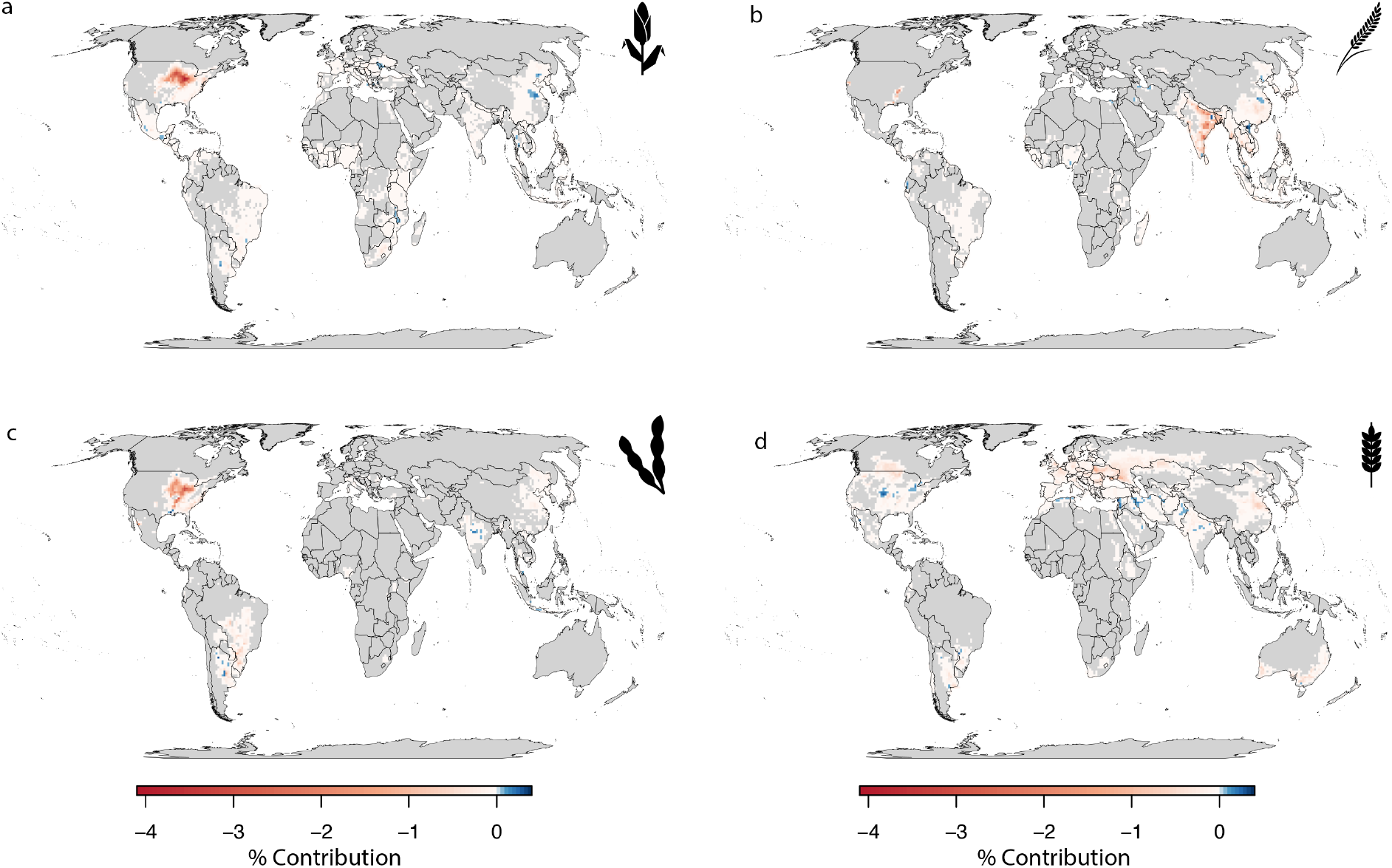
Local contributions to global variance of crop production during 1961-2008. a) Maize, b) Rice, c) Soya, d) Wheat. Each pixel represents a 100 km x 100 km grid cell’s percent contribution to inter-annual global variance in crop production over the last five decades. Negative values show variance inflating (or destabilizing) locations, and positive values show variance deflating (or stabilizing) locations.

### Observed trends in synchrony 1961-2014

Identifying the influence of individual growing locations on the variance of global crop production is useful for spatial prioritization of efforts to increase the resilience of the food system. However, it does not itself indicate if crop production has become more synchronized or more unstable over time. To address this, we draw on recently developed ecological theory (20, 21), to compute three diagnostic metrics of global food system stability, over six distinct 8-year time windows during 1961-2008. These three metrics are global instability (*CV*_*G*_= *σ*_*G*_/*μ*_*G*_), local instability (*CV*_*L*_= *σ*_*L*_/*μ*_*G*_) and synchrony 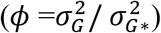, where *σ*_*L*_ is the local standard deviation in production, 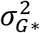, is the global standard variance of production under complete synchronization of local production trends, and *μ*_*G*_ is the mean global production. To maintain an informative picture of the relative severity of losses over the time windows, we compute the numerators of *CV*_*G*_ and *CV*_*L*_ using time detrended production data, and *μ*_*G*_ using observed non-detrended time series. These three quantities are elegantly related such that, 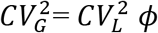: with *φ* acting as a scaling factor that links stability at the local to the global scale (21).

Global instability has shown different trajectories for each of the four major commodities over the past 5 decades (Figure 2). For example, in maize, a massive increase in global instability occurred pre-1984, but has been stabilizing ever since. For soy, global instability peaked during 1961-1968, and has been stabilizing since. Notably, while the planet saw similar crop differences in local instability over 1961-2008 (Figures 2A-B), trends in local instability did not consistently match global instability across crop types. For example, inflections in local instability matched global trends for maize, and to some degree for soy, but this local to global scale trend matching did not occur for wheat and rice (Fig 2A-B). Synchrony peaked for rice, wheat and soy in 1969-77, and for maize in 1977-1984. For maize and soya synchrony declined after this period. For wheat synchrony remained constant in recent decades, and for rice it generally declined over 1977-2000, but rose sharply in 2001-2008 (Figure 2C).

**Figure 2.**
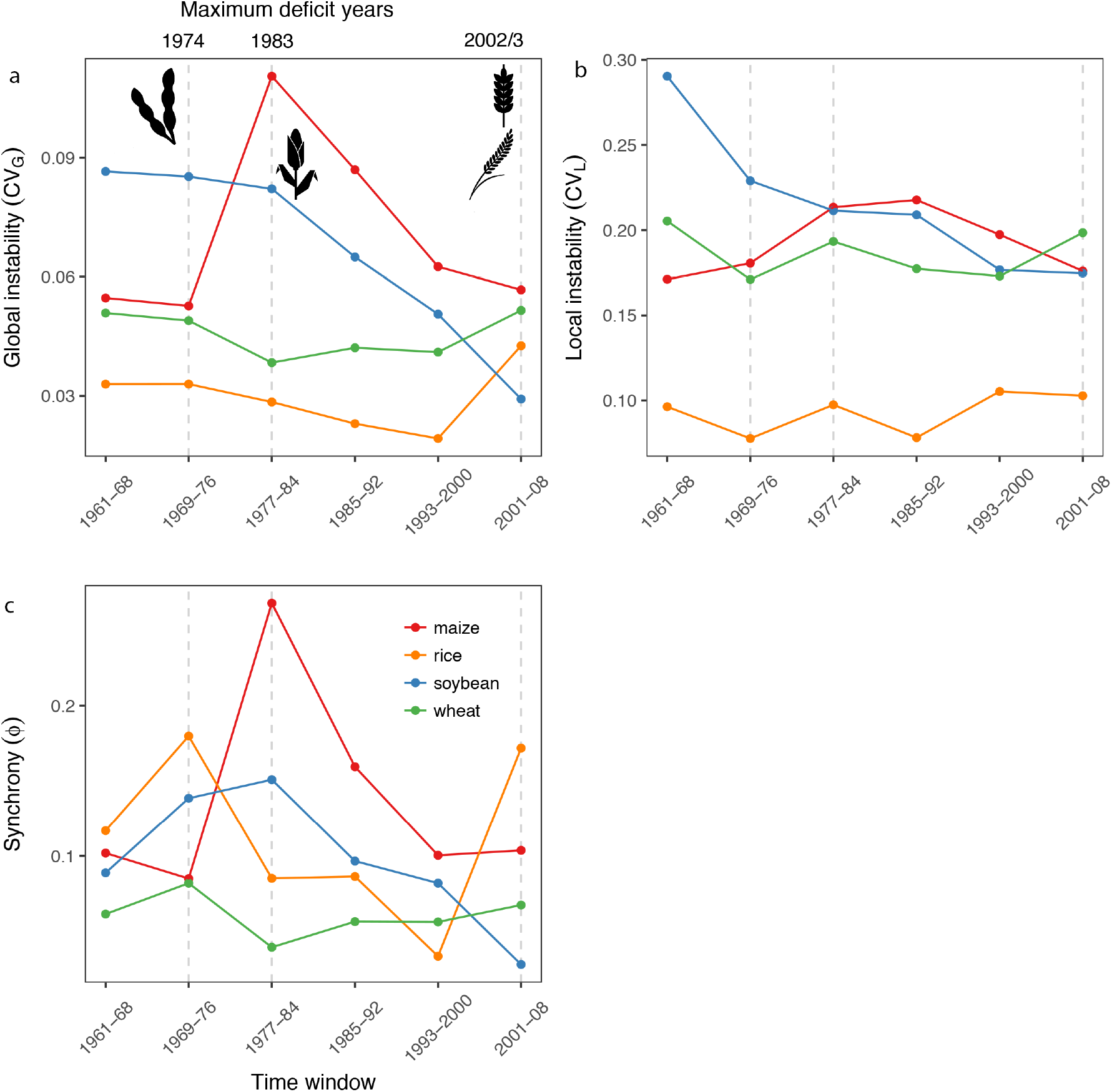
Stability of global crop production during 1961-2008. Trends in global instability result from changes in both local instability, and synchronization in production trends. a) trends in global instability, b) trends in local instability, c) trends in synchrony between local production cells. Dashed gray lines show time windows in which maximum negative deviations from mean production occurred i.e. 1974 for soybean (−15%), 1983 for maize (−23%), 2002 for rice (−8%) and 2003 for wheat (−8%). Synchrony is a unitless metric running from completely synchronous local production trends (1) to completely asynchronous local production trends (0), and scales local to global instability through the following relationship: 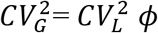.

Notably, synchrony in local production trends matched global instability, and was closely associated with the maximum production deficit in maize (a −23% deviation from the mean trend in 1983) and soy (−15% deficit in 1974) (Figs 2A, 2C). The maximum inter-annual deficit of rice (a −8% loss in 2002) was also associated with the upsurge in synchrony between rice growing regions in 2001-2008. Although maximum losses of −8% in 2003 for wheat were more dependent on increases in local instability than synchrony *per se*, reductions in synchrony played a clear buffering of local destabilization of global wheat production during 1977-84 (Figs 2A-2C).

Taken together, these results indicate that there is little doubt that the degree of synchrony between crop growing locations in the planet has played an important role in regulating the stability and variation in global crop production over the period 1961-2008. Importantly, the food system has, in the general case, trended towards decreased synchronization in local production for maize and soy, which helped stabilize global production trends. Increases in synchronization, where they have occurred, have historically lead to notable destabilizing effects on global crop production.

### Losses under synchronized crop failure

Using the historical data, we constructed a worst-case scenario event under complete synchronization of production trends for each of the four crops, and compared expected losses under this setting to the losses witnessed in the observed trends. We set up our thought experiment to occur in the final year of the dataset, in 2008 (where losses impacts would be closest to the present day due to increasing overall production for all commodities). To estimate the baseline losses, we identified the number of standard deviations that the maximum negative residual from the mean time trend fell over 1961-2008 (−1.8*σ* for soy, −2.9*σ* for maize, −3.6*σ* for rice and −2.3*σ* for wheat), equivalent to the lower bounds of a 100% historical prediction interval for production over the time period. Then to calculate the losses under the worst-case scenario (WCS), we estimated the inflation of the standard deviation in the data under synchrony using the variance-covariance matrix of production trends, and multiplied this by baseline losses for each crop to obtain a maximum negative deviation under synchrony. The baseline losses were −12% for maize, −4% for soy, −8% for wheat, and −8% for rice. Under complete synchrony, the maximum deficits skyrocketed by a factor of three, reaching −36% for maize, −18% for soy, −35% for wheat, and −25% for rice.

Theoretically, there are two types of strategies that could be used to offset the deficits during a worst-case scenario event: mean increasing strategies and variance reducing strategies. Variance reducing strategies, can be implemented by diversifying genotypes, by adapting climate smart cropping systems, by using either ecological engineering, or developing technological infrastructure to resist environmental stressors. Mean increasing strategies on the other hand, can be achieved through expansion of agricultural land, through increasing yield ceilings and decreasing yield gaps. In reality the additional gains from mean increasing strategies would need stocking capacity, but here we are only interested in whether, in principle, the quantities of food generated would be sufficient to offset the losses, and so assume stocking capacity scales with mean production. We assessed the ability of each of these different types of strategies to offset the deficits we observed in our thought experiment of synchronized production, with four radical independent scenarios: (1) “Local variance reduction” = WCS+ 50% reduction in variance in production for every grid cell across the world; (2) “Breadbasket variance reduction” = WCS+ reducing the variance of grid cells in the 90-100th percentile of top producers by 50% percent; (3) “Closing production gaps”= WCS+ increasing production of the bottom 0-50th percentile of producers to 50%; (4) “Raising production ceilings”= WCS+ increasing production of grid cells in the 90-100th percentile of top producers by 50%.

Interestingly, the radical increases in total global production achieved by raising production ceilings or closing production gaps were completely sufficient to offset the −18% to −36% deficits under historical production synchronized failure for maize, rice, wheat and soy (Figs 3 A-D). Mitigation with radical variance reducing strategies on the other hand, were both only able to offset about 10-20% of total losses (Figure 3 A-D). Focusing on reducing variance in breadbaskets performed most poorly for all crops, while the best performing scenario was raising production ceilings. This analysis shows clearly that mean increasing mechanisms are a powerful tool for offsetting the losses under synchronized production failure, and that variance reducing strategies, whilst no-doubt important, will even under radical implementation, be insufficient to tackle the deficits under synchronized crop failure. While it could be argued that the perturbation in each mitigation strategy were arbitrary, they provide a benchmark for what we might expect for different kinds of approaches to stabilize a failing food system.

**Figure 3.**
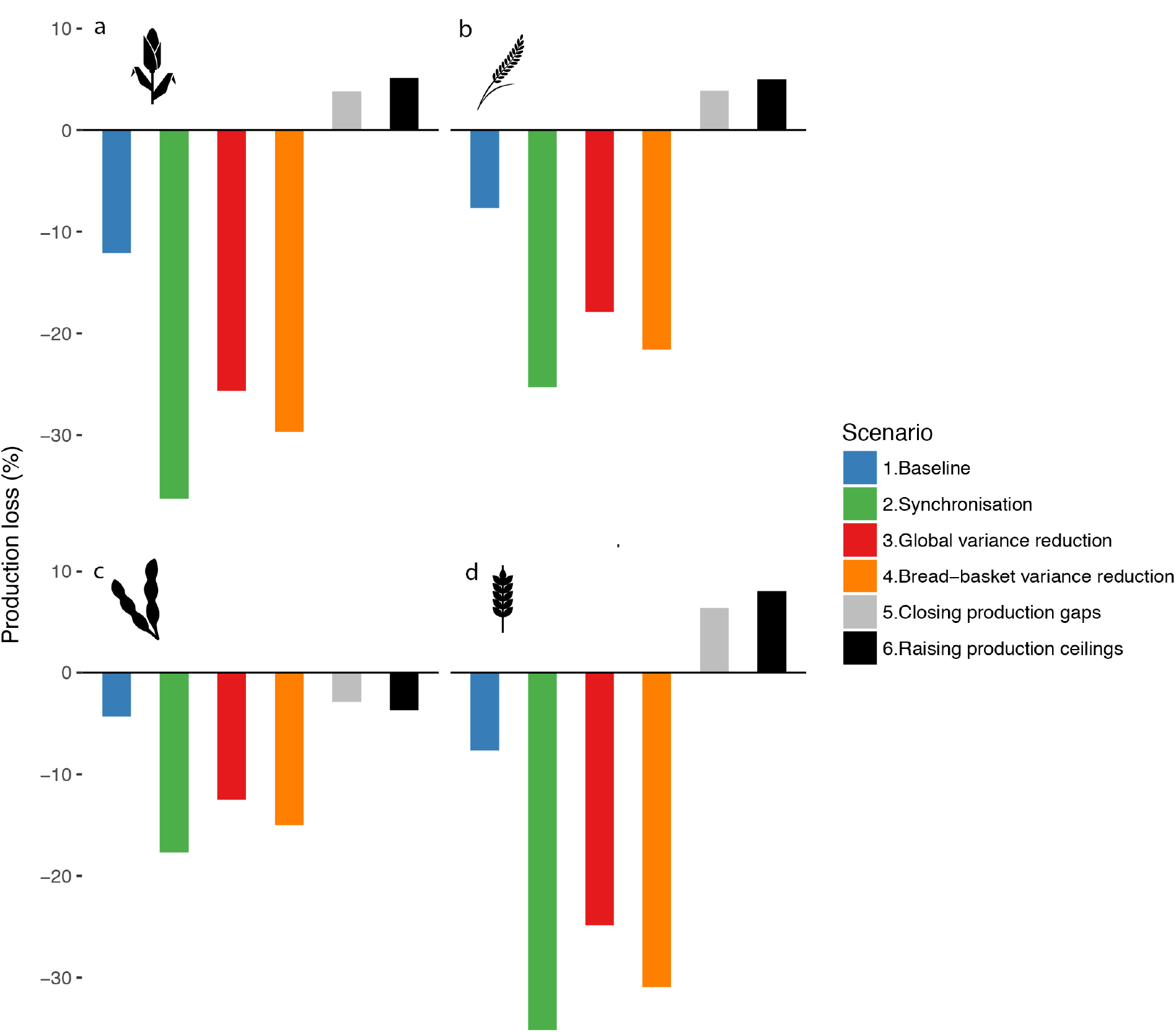
Strategies to cope with losses in a synchronized food system. a) Maize, b) Rice, c) Soya, d) Wheat. Baseline= deviations from the mean for major crop commodities in 2008, estimated using the lower bounds of the distribution of negative deviations from global trends over 1961-2008 (−1.8*σ* for soybean, −2.9*σ* for maize, −3.5*σ* for rice and −2.4*σ* for wheat). Synchronization= worst case scenario (WCS), estimated by inflating baseline deviations by the variance increase due to complete syncrony in local production trends. Local variance reduction = WCS+ 50% reduction in variance in production for every grid cell across the world; Breadbasket variance reduction = WCS+ reducing the variance of grid cells in the 90-100th percentile of top producers by 50% percent; Closing production gaps= WCS+ increasing production of bottom 0-50th percentile of producers by 50%; Raising production ceilings= WCS+ increasing production of grid cells in the 90-100th percentile by 50%.

## Discussion

There are four main take-homes from this analysis. First is that historical records show that for major commodities such as maize and soy, global crop production systems have not tended towards synchronized failure. Second is that typically the discussion of meeting global future food demand has to date been predominantly centered on mean production trends (22–24), with little or no attention to the inter-annual variance in production. Our analysis suggests that the losses under a worst-case scenario could be anything from 18-36% of annual production for all major commodities, which is roughly a quarter to a half of the extra quantities of these crops required to meet projected population increase and consumption demand in 2050 (23, 25). Third, is that our results indicate that variance reducing strategies – such as adapting climate smart cropping systems (26, 27), using either ecological engineering, or developing technological infrastructure to resist environmental stressors (28) – even if widely successful, are may struggle to offset global losses under synchrony. Mean increasing strategies, such a yield gap closure (29) or yield ceiling raising (30), or a mixture of multiple variance reducing and mean increasing strategies should be considered together to mitigate losses. Most importantly, as many cropping systems are likely to exhibit positive mean-variance relationships, the best innovation will be to develop mean increasing strategies that do not inherently raise variances. Fourth, if we are going to use increased production in good years, to deal with losses in the worst years, we need an effective means to food. Importantly, as food demand increases into the future, the requirements for stocks will increase, even if the worst-case scenario is not reached (7). The infrastructure development needed to ensure the future resilience of the global food system is critical.

Our analysis provides an important starting point to begin to quantify the historical risk of synchronized crop failure, and to assess the practical importance of alternative strategies for a more resilient food system in the future. We see three next steps from this work. First, our results indicate that we need a better understanding of how to engineer or maintain asynchrony into the food system wherever possible. There are locations in the world, which are stabilizing global food production, and there is also evidence that the food system has become less synchronized over time. Why this happens, how much of a role climate plays, and how much leverage humanity can have on this aspect of the food system is important to understand. Second, while we addressed crops that make up the vast majority of calorie production on the planet, we only addressed four major crop commodities. This is largely due to data limitations of availability of time series of crops at subnational resolution. The development and availability of global time series data for other important commodities would enable similar analysis for other important crops not included here. Third, we only considered crop production in this work, and expanding our analysis to the stability of components of nutrition (e.g. calories, micronutrients, fats) or to stability of food prices will be an essential next step to better understand the human dimension of resilience in the food system for the future (e.g. 33, 34). Investigating these avenues of research offers a key opportunity to better develop strategies towards a more resilient, safe, and food secure future.

## Materials and Methods

### Dataset

We used globally representative census data on the area and yield of four major commodity crops (rice, maize, wheat, soy), for the years 1961-2008. We computed production (as the product of area x yield) for each producing grid cell in the world and reprojected the data to equal area 100km x 100 km grid cells. Full details of the creation of the original gridded 0.083 degree products are given in earlier publications (18, 33).

### Maps

To create the maps of the local contributions to global variance in production, we first detrended the production time series to ensure the contributions reflect year-to-year variation (which would otherwise be swamped by technology led increases in production over 1961-2008), and then we computed the following index for each focal grid cell on the planet for each crop: 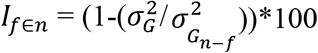, where 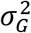 is the global variance and 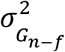 is the global variance when a given grid cell *f* is removed from the total number of producing grid cells *n*. We computed this index independently for each of the four crops used in our analysis prior to mapping.

Formally, 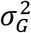 is equal to 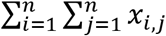, where *i* and *j* are the respective rows and columns of a symmetric variance-covariance matrix *X*. The diagonal elements of *X* represent the temporal variance in production of a given grid cell over 1961-2008, and the off-diagonal elements of *X* represent the temporal covariances in production between grid cells over the same time period. Removing *f* can either increase or decrease 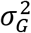. As such, *I*_*f*_ represents the stabilizing or destabilizing effects of grid cell *f* on global crop production over the time period, resulting from inter-annual variation. *I*_*f*_ is in units of percent, where negative values represent a grid cells percent inflation of variance, and positive values represent a percent deflation of variance, of global production over the time period 1961-2008.

### Historical trends

To compute historical trends, we calculated the global instability (*CV*_*G*_), local instability (*CV*_*L*_) and synchrony (*φ*), for the four crops within 8-year windows of the 1961-2008 time period. We draw on recent theory developed for scaling stability in productivity in ecology.

As defined above, for a given set of production time series (i.e. food producing grid cells in the world), the global variance is: 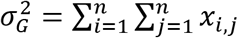, and, the local variance is:

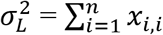 where 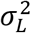 is the sum of all local variance in production for a given crop, and *x*_*i*,*i*_ are the diagonal elements of the symmetric variance-covariance matrix *X*. Importantly, we would expect 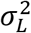 to equal, 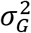, if crop producing regions were uncorrelated with each other, i.e. when all off-diagonal element of *X* equal zero.

The global and local standard deviations are thus: 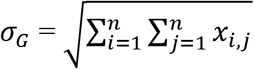, and, 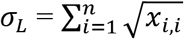 respectively. Using these estimates of global and local variance, we define the global instability and local instability in crop production as the global and local coefficients of variance in production for each of the crops at each time window in the analysis: *CV*_*G*_= *σ*_*G*_/*μ*_*G*_ and, *CV*_*L*_= *σ*_*L*_/*μ*_*G*_ respectively, where *μ*_G_ is the non-deterended mean of global production.

Note, that this formulation (with non-detrended production data for the mean, and time detrended data for the standard deviation) overcomes the influence of non-stationarity in the mean on interannual variance (i.e. due to technology change), but ensures an informative picture of the relative severity of losses is maintained (e.g. a −50% deviation from the mean in 1961 is much smaller in absolute terms than a −50% deviation in 2008).

Finally, we computed the third diagnostic metric, synchrony: 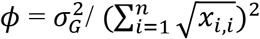, where *φ* is the synchrony between the all the producing grid cells in the world for a given crop. The denominator of this ratio, 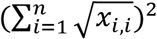, or 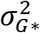, is equal to 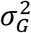 when all elements of the correlation matrix of producing grid cells (*P*) have correlation of *ρ* = 1. This index is bounded by 1, complete synchrony and approaches 0, when all the elements of *P* tend from 0 to −1, to give complete asynchrony. This metric is useful because it shows how close we have been globally to the ‘worst case’ scenario of complete synchronous production dynamics over the period 1961-2008.

Global instability (*CV*_*G*_), local instability (*CV*_*L*_) and synchrony (*φ*) are related such that: 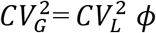. With *φ* acting as a scaling factor that links stability at the local to the global scale

### Scenario planning

Using the historical data, we constructed a worst-case scenario event under complete synchronization of production trends for each of the four crops, and compared expected losses under this setting to the losses expected under the observed trends. We set up our thought experiment to occur in the final year of the dataset, in 2008. To estimate the baseline losses, we used the number of standard deviations that the maximum losses fell over 1961-2008 (−1.8*σ* for soy, −2.9*σ* for maize, −3.6*σ* for rice and −2.4 *σ* for wheat), to gain the lower bounds of a 100% historical prediction interval for production of this period. To estimate the losses under the worst-case scenario (WCS), we estimated the inflation of the standard deviation in the data under synchrony using the variance-covariance matrix of production trends, i.e. 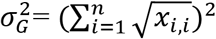, and multiplied this by the baseline deviations for each crop to obtain a maximum negative deviation from the mean under synchrony.

We then ran four mitigation scenarios under the ‘worst case scenario’: (1) “Local variance reduction” = WCS+ 50% reduction in variance in production for every grid cell across the world; (2) “Breadbasket variance reduction” = WCS+ reducing the variance of grid cells in the 90-100th percentile of top producers by 50% percent; (3) “Closing production gaps”= WCS+ increasing production of bottom 0-50th percentile of producers by 50%; (4) “Raising production ceilings”= WCS+ increasing production of grid cells in the 90-100th percentile by 50%. And then determined how much each of these strategies was able to offset production deficits under complete synchrony, making use of the fact that the standard deviation of global crop production, *σ*_*G*_ under complete synchrony, when all elements of *P* = 1, is simply, 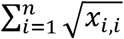, which is equal to the baseline worse case, and otherwise *σ*_*G*_ is equal to 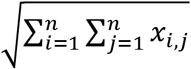.

A full set of reproducible **R** (34) script is supplied as Supplementary Information to undertake the entirety of the analysis presented in this paper.

## Author contributions

ZM designed the research, ran the analysis, and wrote the paper, with ongoing input from NR.

## Acknowledgements

Many thanks to Deepak Ray for sharing the data set. NR and ZM were funded by NSERC and Genome BC.

## Data availability statement

Full details of the data used in this paper are given in the Supplementary Information.

