## Supplementary Materials for "On the synchronized failure of global crop production"

### On the syncrhonized failure of global crop production
### Mehrabi and Ramankutty 2018

March 13, 2018

### 1 Aim of document

The aim of this document is to provide the Supplementary Information for Mehrabi and Ramankutty 2018. This Supplementary Document was created with `knitr` [6], a software that combines the typesetting system LaTeX and the statistical computing language **R** [2], and allows for full reproduction of our results.

This document is split into three sections. In section 2 we illustrate how we make our analysis fully reproducible. Section 3 explains a bit of background to the data we are working with and shows how we manipulated the data to get it ready for analysis. Finally, section 4 walks step by step through the analysis that underpins the results presented in our paper.

### 2 Checkpoint

Here we call the **R** package `checkpoint` [1]. This will create a local library on your computer and install a copy of the packages required by this project as they existed on CRAN on the specified snapshot date, and update the **R** session to use these packages. This helps make our analysis fully reproducible on your machine.

```
require(checkpoint)
checkpoint(snapshotDate = "2018-02-01")
```

Note that the **R** version used here is 3.4.2 (2017-09-28). Using other versions of **R** should not have any influence on the results obtained. The function call just above installed all packages used in this document, as available on Febuary 1st 2018 in the "home" directory of my computer.

### 3 Data preparation

#### 3.1 Data set overview

The data we will be using for this analysis is Deepak Ray and colleagues data, which was described in detail in [4] and [3]. Briefly, these are globally representative, census data on the area, yield, of four major commodity crops (rice, maize, wheat, soybean), for the years 1961-2008. Deepak Ray and others [3] gridded this census data across the earths surface to 0.5 degree tiles, and we will be using this gridded product for the analysis in this paper.

Deepak Ray and colleagues data is not in the correct format for the analysis we wish to perform. First the data is in netcdf (.nc) format, and housed in separate years, and we need to import this into **R** and analyse it as a seamless time series. Second, the data is presented as separate Area and Yield files for each crop, and we will need to compute Production. Third, we wish to scale the data from its current spatial scale of c. $10km^2$ at the equator to (i.e. 0.083 degrees) to c. $100km^2$ (i.e. 1 degree). And finally, as we want to work with tiles of comparable spatial extent, we need to reproject the data to an equal area grid. Each of these steps are outlined below. Most of this data preparation is not evaluated in this document, although this can be easily switched by changing the 'evaluate=FALSE' argument in the `knitr` headers of the code sections to 'evaluate=TRUE'.

#### 3.2 Getdata function

Below I've written a function to read in the netcdf data from multiple files and create a common data frame containing all of the data in them. Deepak Ray's data has the same lat long referencing and time periods for all datasets, which makes this step of manipulation quite easy.

```
getdata<-function(path="", pat="") {
setwd(path)
file.list = list.files(pattern=pat)
files = lapply(file.list, nc_open) #get all the files
All=NULL
for (i in 1:length(files)) {
  Data<- as.vector(ncvar_get(files[[i]], "Data")) #pull in data variable
  All<-cbind(All, Data) #bind data variables together

}
colnames(All)<-file.list #assign data nc file source info
lapply(files, nc_close) #close nc files
All
}
```

#### 3.3 Create production variable

Below is the code to create a production estimate based on Fractional area (percent under cultivation) and yield (tonnes/ha). Some crops have multiple cropping seasons, which is why their F. Area is greater than 1. We have set crops to a maximum of two cropping seasons per year (e.g. F. area of 2), except rice which is set to three. Fractional area's less than 0.05 percent are removed. We took these two steps to ensure only sensible area values included. Finally production files for each crop are saved to their own `.rds` file.

```
library(ncdf4)
library(raster)
library(plyr)

#set out the paths for crops.
crops<-list("soybean", "maize", "rice", "wheat")
folder<-"/Users/pumpkinjr/Sync/academic/projects/UBCload/Data/Deepak's historical data/" #path to raw data
newfolder<-"/Users/pumpkinjr/Sync/academic/projects/UBCload/Data/Deepak's historical data/processed/" #new folder for output

for(i in 1:length(crops)) {
 path=paste(folder,crops[i], sep = "")
 Area<-getdata(path=path, pat="*Area_ver8.nc") #this is in tonnes/ha
 Yield<-getdata(path=path, pat="*Yield_ver8.nc") #in % area under cultivation in grid square

#make sure % area <0.05 and >2 are not present in the data, these values make little sense
Area[Area<0.05]<-NA
ifelse(path=="/Users/ziamehrabi/Documents/Data/Deepak's historical data/rice",Area[which(Area>3)]<-NA,
       Area[which(Area>2)]<-NA) #rice has 3 growing seasons

#estimate area of grid cells.
 test<-raster("/Users/ziamehrabi/Documents/Data/Deepak's historical data/soybean/Soybean_1961_Area_ver8.nc") #template
 area<-area(test)
 ar.ha<-(values(area)*100) #convert from km2 to ha
 Prod<-(Area*Yield*ar.ha*1000) #get production per cell, in kg

#save file to RDS
path2=paste(newfolder,crops[i], sep = "")
saveRDS(object = Prod, file = paste(path2, "production.rds", sep = ""))

#remove files to free up ram
rm(Area)
rm(Yield)
rm(Prod)
}
```

#### 3.4 Upscaling the data

Next we upscale the data, and reproject it into Eckert's Equal Area. The final product is production for each crop in $100km^2$ grid cells across the earth, for the years 1961-2008. These are the units of analysis for the manuscript. The code below runs for each input `.rds` file created in the preceding section, and creates a new `.rds` file with the aggregated and reprojected data.

```
rm(list = ls())#clear space
library(ncdf4) #load libaries
library(raster)
library(plyr)

newfolder<-"/Users/pumpkinjr/Sync/academic/projects/UBCload/Data/Deepak's historical data/processed" #folder for output

crops<-list("soybean", "maize", "rice", "wheat")

for(i in 1:length(crops)) {
  crop<-crops[i]
   path=(paste(crop,"production.rds", sep = ""))
   Prod <- readRDS(path)

#read back in production file to get co-ordinates and append
 names<-colnames(Prod)
  test2<-nc_open("/Users/pumpkinjr/Sync/academic/projects/UBCload/Data/Deepak's historical data/soybean/Soybean_1961_Area_ver8.n
  latlon<- expand.grid(lon=ncvar_get(test2, "longitude"),lat=ncvar_get(test2, "latitude")) #get latlong of original
  Prod<-cbind(latlon,  Prod) #bind latlon to production dataset
  nc_close(test2)

#create a spatial data frame of the original data (for converting to raster) and raster for aggregating up cells.
coordinates(Prod) <- ~ lat+ lon #make spatial
rasterD <-raster(ncol=4320, nrow=2160) #set original extent and dimensions. raster() assigns wgs84 as default ell+ datum

#aggregate then reproject (quicker, less accurate).
 v=list(ncol(Prod))
 for(i in 1:ncol(Prod)) {
   values(rasterD)<-Prod[[i]]

    values(rasterD)<-ifelse(is.na(values(rasterD)), 0,values(rasterD) )#setting NA to zero (required)
    Raster.ag <- aggregate(rasterD, fact=12, fun=sum, na.rm=T ) #aggregate to 300km2 grid
    Raster.D.equal <- projectRaster(Raster.ag, res=c(100000, 100000),
    crs="+proj=eck4 +datum=WGS84 +ellps=WGS84 +towgs84=0,0,0",
    method="ngb", over = T) #convert to equal area

values(Raster.D.equal)<-ifelse(values(Raster.D.equal)== 0,NA, values(Raster.D.equal) )#reset zeros to NA.

   v[[i]]<-values(Raster.D.equal)
 }

 #add col names and convert back to df
 v.df<-data.frame(matrix(unlist(v), ncol=48))
 colnames(v.df)<-names
 scaled.prod<-cbind(v.df, coordinates(Raster.D.equal))
  saveRDS(object = scaled.prod, file = paste(crop, "production100km2.rds", sep = ""))
}
```

### 4 Analysis

In this section we perform the analysis presented in the manuscript. It is split into 3 general subsections. Firstly, we map out the local contributions to global variation in crop production for the time period of our study. Then we break the time series down to assess how local stability or synchrony has changed over time, and how much each has contributed to global instability for the four crops used in our analysis over the period 1961-2008. Finally we undertake a thought experiment to see how large production deficits would have been in 2008, if all production cells across the world were synchronized. We then see what how much of an effect different mitigation strategies have on offsetting production loss.

#### 4.1 Estimate local contributions

##### 4.1.1 Variance contribution index

In this subsection I map out the local contributions to global variance of crop production for four major crop types. These local contributions are due to both local variation in production at the grid cell level, and the correlation in production between different grid cells across the world. First we detrend the production time series (to ensure the contributions reflect year-year variation, which would otherwise be swamped by mean technology lead increases in production over 1961-2008), then we compute the following index for each focal grid cell on the planet for each crop:

$I_{f \in n}$ = (1-($\sigma_G^2$/ ${\sigma_G}_{n-f}^2$))*100

Where $\sigma_G^2$ is the global variance and ${\sigma_G}_{n-f}^2$ is the global variance when a given grid cell $f$ is removed from the total number of producing grid cells $n$. We computed this index independently for each of the four crops used in our analysis.

Formally, $\sigma_G^2$ is equal to $\sum_{i=1}^{n}\sum_{j=1}^{n} x_{i,j}$ , where $i$ and $j$ are the respective rows and columns of a symmetric variance-covariance matrix $X$. The diagonal elements of $X$ represent the temporal variance in production for a given grid cell over 1961-2008, and the off-diagonal elements of of $X$ represent the temporal covariances in production between grid cells over the same time period. Removing $f$ can either increase or decrease $\sigma_G^2$. As such, $I_f$ represents the stabilizing or destabilizing effects of grid cell $f$ on global crop production over the time period, resulting from inter-annual variation. $I_f$ is in units of percent, where negative values represent a grid cells percent inflation of variance, and positive values represent a percent deflation of variance, of global production over the time period 1961-2008.

In the code that follows we first detrend the time series of production, then compute $I_f$ for each $n$ grid cell. We do this independently for each crop. The output is stored as a list named "all.crop.var". Each element of the list is a crop, and the numerical values in that list element represent the destabilization index. Note that the co-ordinates of the grid cell have been removed from this final product, we will need to add them in back later.

```
rm(list = ls())#clear space
```

```r
library(raster) #load libraries
library(sp)
library(maptools)
library(grid)
library(rgdal)

crops<-list("maize" , "rice","soybean","wheat" )

all.crop.var=list()

for(i in 1:length(crops)) {
  crop<-crops[i] #assign crop
   path=(paste(crop,"production100km2.rds", sep = "")) #create path
   w <- readRDS(path)  #read in
D<-w[ , -which(names(w) %in% c("y","x"))] #remove co-ords.

# make all NA's zero
D[is.na(D)] <- 0 # replace NA with zeros, otherwise the "Dn" variable will include NAs in denominator of global produciton ratio.
c=colSums(D, na.rm=T) #global production trends for whole dataset
Dn<-t(apply(D,(1),function(x) c-x)) #get differences (global production minus contribution of grid cell), transpose out.

#get variance of detrended production for data with grid cells removed
Dn2<-matrix(nrow=nrow(Dn),ncol=ncol(Dn)-1) #make matrix for detrended
for(i in 1:nrow(Dn))
{
Dn2[i,]<-diff(Dn[i,], lag=1, differences=1) #detrend each row
}
var.detrend<-apply(Dn2,1, var) #get variance of each row

var.dc<-var(diff(c, lag = 1, differences = 1)) #get variance of mean deterended production

#get ratio, >1 then detsabilizing, < 1 then stabilizing
#is global variance more or less than with the cell?
v.ratio<-var.dc/var.detrend

#remove 1's cases where grid cell prod is 0 and so the ratio is exactly equal to 1.
v.ratio.new<-replace(v.ratio, v.ratio==1, NA)
v.ratio.new<-(round((1-v.ratio.new)*100, digits=1)) #change to percentage

#bind alltogether in dataframe + save
name <- crop[[1]]
all.crop.var[[name]] <- v.ratio.new
}
```

##### 4.1.2 Summary of the local contributions

In this section we quickly summarise the distributions of the variance contribution index for each crop. Expanding this summary we will see the maximum variance inflating and deflating locations differ for each crop and so does the mean value. Most locations are all close to zero in their values, which means they have no practically significant impact on year-year variation in global crop production.

```r
summarydat<-lapply(all.crop.var, function(x) quantile(x, probs = seq(0, 1, 0.01),na.rm = T))
```

##### 4.1.3 Plot local contributions

Next we plot the local contributions to global variance for each crop. We will use the co-ordinates called in the loop in the previous last section to create a common plotting space with the correct dimensions, projection and co-ordinate system. Once we have this we will assign the data to this common template, and plot out the data on global maps.

```
data<-data.frame(w$x, w$y, v.ratio.new)  #fill results into a matrix
coordinates(data) <- ~ w.x+ w.y
gridded(data) <- TRUE
raster.w<-raster(data)
raster.x <-raster(res=c(100000, 100000), crs="+proj=eck4 +datum=WGS84 +ellps=WGS84 +towgs84=0,0,0")
projection(raster.w) = projection(raster.x)
```

Finally, we plot out the maps. First we set up the color schemes, and the min and maximum values for the common legend for plotting, and the breaks for the colors.

```
#get min and max across the four plots for common legend
min_ = min(c(all.crop.var[[1]], all.crop.var[[2]], all.crop.var[[3]], all.crop.var[[3]]), na.rm=T)
max_ = max(c(all.crop.var[[1]], all.crop.var[[2]], all.crop.var[[3]], all.crop.var[[3]]), na.rm=T)

#assign breaks
breaks<-c(seq(min_,0, length.out=100))
breaks2 <-c(seq(0.0001,max_, length.out=100))

##blue red
color.palette= rev(colorRampPalette(c("#053061", "#2166ac", "#4393c3","#f7f7f7"))(length(breaks2)))  ##with blue
color.palette2= rev(colorRampPalette(c( "#fffbf9",  "#f4a582","#d6604d","#b2182b"))(length(breaks))) ##red side

#join them together
breaks.all<-c(breaks2, breaks)
colors.all<-c(color.palette2,color.palette)
```

Then we run a loop to plot each crop in turn. We first set the plotting frame in to be 2x2 units using the graphical parameters function (par()), and assign each element of the “all.crop.var” list to the plotting frame. Some manipulation of the plotting area using par() was undertaken to ensure the plot is compact for publication. We run the loop for the first and second rows of the panel, with the legend included in the second row to make the presentation neater.

```
data(wrld_simpl) #get world map data

par(mfrow = c(2,2),#set up plottting area
        oma = c(26,0,0,0) + 0.1,
          mar = c(0,0,0,0) + 0.1)

for(i in 1:2){
values(raster.w)<-all.crop.var[[i]]#assign values to raster
wrld <- spTransform(wrld_simpl, crs(raster.x))
plot(wrld, ylim=c(-99910,99820),  col ="light gray",border=NA, bg="white" )
plot(raster.w,col = colors.all, cex=1, breaks=breaks.all, add=TRUE, legend=F)
plot(wrld, ylim=c(-99910,99820), col='transparent', border="black", add=TRUE,lwd=0.2 ) #add transparent borders
}

for(i in 3:4){
values(raster.w)<-all.crop.var[[i]]#assign values to raster
wrld <- spTransform(wrld_simpl, crs(raster.x))#plot out
plot(wrld, ylim=c(-99910,99820),  col ="light gray",border=NA, bg="white" )
plot(raster.w,col = colors.all, cex=1, breaks=breaks.all, add=TRUE, horizontal=TRUE,
     legend.mar =0, legend.width = 1.1, legend.shrink=0.55,legend.args=list(text='',
                     font=2, line=1, cex=1), axis.args=list(at=pretty(c(round(min_, digits=2),0,round(max_, digits=2))),
                   labels=pretty(c(round(min_, digits=2),0,round(max_, digits=2)))))
plot(wrld, ylim=c(-99910,99820), col='transparent', border="black", add=TRUE,lwd=0.2 ) #add transparent borders
}
```

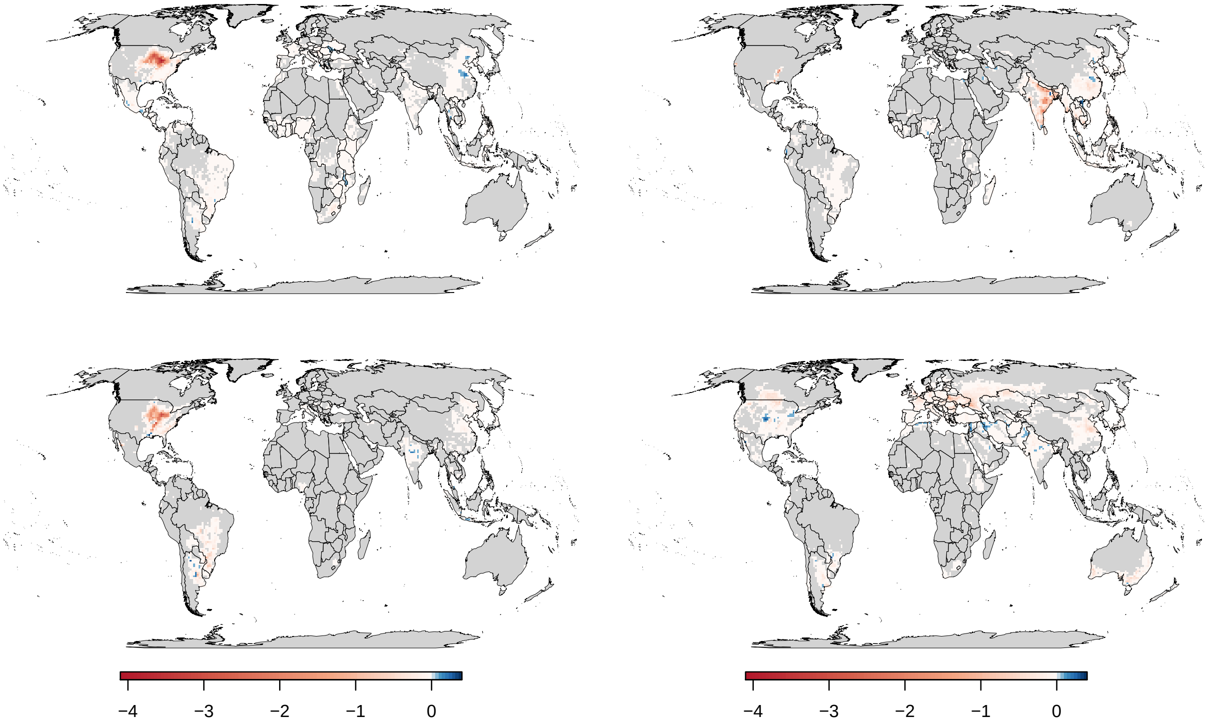

Figure 1: Local contributions to global variance of crop production over 1961-2008. Each pixel represents a 100 km x 100 km grid cells percent contribution to inter-annual global variance in crop production for the last five decades. Negative values show variance inflating locations, and positive values show variance deflating locations. Crops shown, from top right to bottom left: Maize, Rice, Soya, Wheat

#### 4.2 Decadal analysis

##### 4.2.1 Summary indices

In this subsection we calculate the global instability ($CV_G$), local instability ($CV_L$) and synchrony ($\phi$), for the four crops within 8 year windows of the 1961-2008 time period. We draw on recent theory developed for scaling stability in productivity in ecology [5]. First we detrend the data (using loess regression to account for non-linearities in production trends), then calculate these summary statistics, before storing them in a common dataframe "all.crop" ready for plotting.

As defined above, for a given set of production time series (i.e. food producing grid cells in the world), the global variance is:

$\sigma_G^2 = \sum_{i=1}^{n} \sum_{j=1}^{n} x_{i,j}$

And, the local variance is:

$\sigma_L^2 = \sum_{i=1}^{n} x_{i,i}$

Where $\sigma_L^2$ is the sum of all local variance in production for a given crop, and $x_{i,i}$ are the diagonal elements of the symmetric variance-covariance matrix $X$. Importantly, we would expect $\sigma_L^2$ to equal, $\sigma_G^2$, if crop producing regions were uncorrelated with each other, i.e. when all off-diagonal element of $X$ equal zero. [1]

The global and local standard deviations are thus:

$\sigma_G = \sqrt{\sum_{i=1}^{n} \sum_{j=1}^{n} x_{i,j}}$.

and,

$\sigma_L = \sum_{i=1}^{n} \sqrt{x_{i,i}}$

Using these estimates of global and local variance, we define the global instability and local instability in crop productivity as the global and local coefficients of variation in production for each of the crops at each time window in the analysis:

$CV_G = \sigma_G / \mu_G$

and,

$CV_L = \sigma_L / \mu_G$

Where $\mu_G$ is the non-deterended mean of global production. Note, that this formulation (with non-detrended productivity data for the mean, and mean detrended data for the standard deviation) overcomes the influence of non-stationarity in the mean on inter-annual variance (i.e. due to technology change), but ensures an informative picture of the relative severity of losses is maintained (e.g. a -50% deviation from the mean in 1961 is much smaller in absolute terms than a -50% deviation in 2008).

Finally we compute the third diagnostic metric, synchrony:

$\phi = \sigma_G^2 / \ (\sum_{i=1}^{n} \sqrt{x_{i,i}})^2$

Where $\phi$ is the synchrony between the all the producing grid cells in the world for a given crop. This denominator of this ratio, $(\sum_{i=1}^{n} \sqrt{x_{i,i}})^2$ is equal to $\sigma_G^2$ when all elements of the correlation matrix of producing

[1] Indeed, the only difference between $\sigma_G^2$ and $\sigma_L^2$ is that $\sigma_G^2$ includes the off diagonals of $X$. These off diagonal elements can be obtained, for any given set of correlations, from the following matrix: $M = \sqrt{X_{diag}} * \sqrt{X_{diag}}^T * P$,i.e where $M$ is equal to the outer product of the vector of standard deviations from each producing grid cell multiplied by the specified correlation matrix ($\rho$ for each pair of crop producing cells). When all $\rho$ in $P = 0$, then the variance of the sum is equal to the sum of the variances, $\sigma_G^2$ will equal $\sigma_L^2$, and we will have direct mapping of local and global instability. When all $\rho$ in $P$ are 1, then the sum of the off diagonal elements of $M$ will be exactly equal to $(\sum_{i=1}^{n} \sqrt{x_{i,i}})^2 - \sum_{i=1}^{n} x_{i,i}$, and global instability will be magnified relative to local instability. As $\rho$ in $P$ approach -1, then the sum of the off diagonal elements of $M$ will begin to cancel the sum of the diagonal elements of $M$, and global instability will be much less than local instability (note, this may occur prior to average $\rho$ values attaining 0).

grid cells ($P$) have correlation of $\rho = 1$. This index is bounded by 1, complete synchrony and approaches 0, when all the elements of $P$ tend from 0 to -1, to give complete asynchrony.This metric is useful because it shows how close we are globally we have been the 'worst case' scenario of complete synchronous production dynamics over the period 1961-2008.

These three quantities are related such that:

$$CV_G{}^2 = CV_L{}^2 \cdot \phi$$

With $\phi$ acting as a scaling factor that links stability at the local to the global scale [5]

```r
rm(list = ls())#clear space

crops<-list("maize", "rice","soybean", "wheat")
all.crop=data.frame()

for(i in 1:length(crops)) {
  crop<-crops[i] #assign crop
   path=(paste(crop,"production100km2.rds", sep = "")) #create path
   w <- readRDS(path)  #read in
   w<- subset(w,, -c(x, y)) #remove co-ordinates

w.compl<-w[rowSums(is.na(w))!=ncol(w),] #remove complete cases of NA
w.compl[is.na(w.compl)] <- 0 #replace NA with zeros

year<-1961:2008
w.compl<-w.compl/10e9 #convert kg to into mega tonnes

w.compl.dec<-list(w.compl[1:8], w.compl[9:16], w.compl[17:24], w.compl[25:32], w.compl[33:40],w.compl[41:48])

w.compl.dec.sum<-lapply(w.compl.dec,function(x) colSums(x))

detrend.w<-data.frame(t(apply(w.compl, 1,function(x) resid(loess(x~year))))) #use loess for detrending

det.w.dec<-list(detrend.w[1:8], detrend.w[9:16],
                detrend.w[17:24], detrend.w[25:32],
                detrend.w[33:40],detrend.w[41:48])  #split data into windows

det.w.dec<-lapply(det.w.dec, t) #transpose to get the grid cells on the cols

diff.cov<- lapply(lapply(det.w.dec,data.frame), cov) #make covariance matrix of the grid cells.
diag.cov<-lapply(diff.cov, diag) #get diagonals of covariance matrix = local variance

sqrt.diag.cov<-lapply(diag.cov, sqrt) #sqrt of diags of covariance matrix = list of local sds

diff.var.glob<-unlist(lapply(diff.cov, sum)) #sum the covariance matrix. = global variance
diff.var.local<-unlist(lapply(diag.cov, sum)) #sum the diagonals = local variance

diff.sd.global<-sqrt(diff.var.glob) #global sd
diff.sd.local<-unlist(lapply(sqrt.diag.cov, sum)) #local sd

diff.sync <- diff.var.glob /(diff.sd.local^2) #synchrony

mean.dec<-unlist(lapply(w.compl.dec.sum, mean)) #non-detrended global means

CVl<- diff.sd.local/mean.dec #local instability
CVg<-diff.sd.global/mean.dec #global instability

#bind all together in dataframe + save
 vars<-c(rep("global variance",6), rep("local variance",6), rep("synchrony",6) , rep("CVl",6) , rep("CVg",6))
 data<-c(diff.var.glob, diff.var.local, diff.sync,CVl,CVg)
 year<-c(rep(1:6, 5))
 cropid<-c(rep(unlist(crop),30 ))
 all<-data.frame(vars,data, year, cropid) #crop specific
 all.crop<-rbind(all.crop, all)
}
```

##### 4.2.2 Plot decadal analysis

In this subsection we plot the data from the decadal analysis above.

```
library(ggplot2) #load libraries
library(gridExtra)
library(cowplot)

all.c.split<-split(all.crop, all.crop$vars) #split buy variable to plot individsually

global<-ggplot(all.c.split$CVg, aes(year,(data), colour=cropid))+
   geom_vline(xintercept = 2, lty=2, color="light gray")+
  geom_vline(xintercept = 3, lty=2, color="light gray")+
  geom_vline(xintercept = 6, lty=2, color="light gray")+
 geom_point()+
  geom_line()+
 theme_bw()+
scale_color_manual(values = c("#e41a1c","#ff7f00", "#377eb8",  "#4daf4a"))+
  ylab(expression(Global~instability~(CV[G])))+
 theme(panel.grid.major = element_blank(), panel.grid.minor = element_blank(),
        legend.position = "none",legend.key = element_blank(),
         legend.key.size = unit(0.5, "cm"), legend.title=element_blank(),
       axis.text.x = element_text(angle=45, vjust=0.5))+
    xlab("")+
scale_x_continuous(breaks=1:6, labels=c("1" = "1961-68", "2" = "1969-76",
                                        "3" = "1977-84", "4" = "1985-92",
                                        "5" = "1993-2000", "6" = "2001-08"))

local<-ggplot(all.c.split$CVl, aes(year,(data), colour=cropid))+
   geom_vline(xintercept = 2, lty=2, color="light gray")+
  geom_vline(xintercept = 3, lty=2, color="light gray")+
  geom_vline(xintercept = 6, lty=2, color="light gray")+
 geom_point()+
  geom_line()+
 theme_bw()+
scale_color_manual(values = c("#e41a1c","#ff7f00", "#377eb8",  "#4daf4a"))+
  ylab(expression(Local~instability~(CV[L])))+
  theme(panel.grid.major = element_blank(), panel.grid.minor = element_blank(),
        legend.position = "none",legend.key = element_blank(),
  axis.text.x = element_text(angle=45, vjust=0.5))+
   xlab("")+
scale_x_continuous(breaks=1:6, labels=c("1" = "1961-68", "2" = "1969-76",
                                        "3" = "1977-84", "4" = "1985-92",
                                        "5" = "1993-2000", "6" = "2001-08"))

sync<-ggplot(all.c.split$synchrony, aes(year,(data), colour=cropid))+
  geom_vline(xintercept = 2, lty=2, color="light gray")+
  geom_vline(xintercept = 3, lty=2, color="light gray")+
  geom_vline(xintercept = 6, lty=2, color="light gray")+
 geom_point()+
  geom_line()+
 theme_bw()+
scale_color_manual(values = c("#e41a1c","#ff7f00", "#377eb8",  "#4daf4a"))+
  ylab(expression(Synchrony~(phi)))+
theme(panel.grid.major = element_blank(), panel.grid.minor = element_blank(),
        legend.key = element_blank(), axis.text.x = element_text(angle=45, vjust=0.5),
      legend.key.size = unit(0.5, "cm"), legend.title=element_blank(), legend.position = c(.8, .8))+
    xlab("Time window")+
scale_x_continuous(breaks=1:6, labels=c("1" = "1961-68", "2" = "1969-76",
                                        "3" = "1977-84", "4" = "1985-92",
                                        "5" = "1993-2000", "6" = "2001-08"))

global<-plot_grid( global,local,sync, align =c("v", "h"), ncol = 2)

global
```

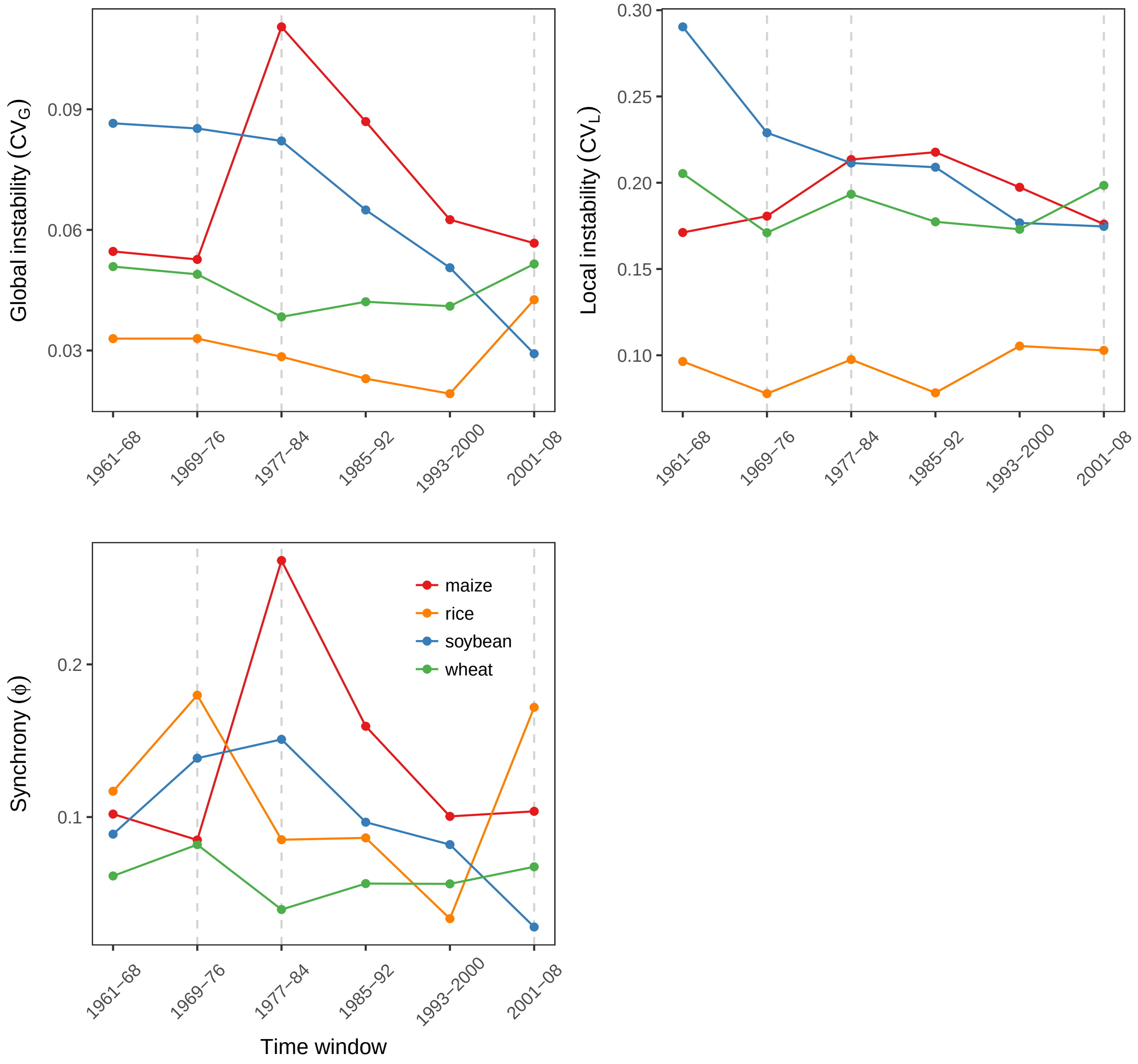

Figure 2: Stability of global crop production 1961-2008. Trends in global instability result from changes in both local instability, and synchronation in production trends. Dashed gray lines show time windows in which maximum negative deviations from mean production occured i.e. 1974 for soybean (-15%), 1983 for maize (-23%), 2002 for rice (-8%) and 2003 for wheat (-8%). Synchrony is a unitless metric running from completely synchronous local production trends (1) to completely assyncronous local production trends (0), and scales local to global instability through the following relationship: $\mathrm{CV}_G^2 = CV_L^2 \cdot \phi$.

#### 4.3 Scenario planning

In this section we run an analysis to understand what the impact of synchronisation would be on the maximum deficits (percent negative deviations from the mean) observed for each of the major crops over the period of 1961-2008. This is a very simple approach that starts by converting residual production values into percentages of the mean trend. After this, we then compute the number of standard deviations that each of these deficits fall by computing the standard deviation of the residuals and dividing each residual value by that standard deviation. In the next step, the standard deviation of the time series of each of the crops under synchrony is determined. And then, to identify the residual under synchrony, the standard deviation of the syncrhonised time series are multiplied by the standard deviation distance that a deficit falls in the years of the maximum deficits. Finally these new residuals under synchrony are converted into percentages based on the non-deterended production time series.

##### 4.3.1 Pre-liminary summaries

Here we identify the year with the maximum percent loss in production, the value of this deviation, and the number of standard deviations from the mean that this occured. As we can see from the table the worst years occured midway in the time series for maize (1983) and soybean (1974), and in the last decade of the time series for rice (2002) and wheat (2003). The observed largest percent losses from the mean trend are of -23% for maize, -15% for soybean, and -8% for both wheat, and rice. In total the worst years represented deviations of -1.8$\sigma$ for soybean, -2.9$\sigma$ for maize, -3.6$\sigma$ for rice and -2.3$\sigma$ for wheat

```r
rm(list = ls())
crops<-list("soybean", "maize", "rice", "wheat")
all.crop.final=data.frame()

for(i in 1:length(crops)) {
  crop<-crops[i] #assign crop
   path=(paste(crop,"production100km2.rds", sep = "")) #create path
   w <- readRDS(path)  #read in
   w<- subset(w,, -c(x, y)) #remove co-ordinates

w.compl<-w[rowSums(is.na(w))!=ncol(w),] #remove complete cases of NA
w.compl[is.na(w.compl)] <- 0 # replace NA with zeros
w.compl.nd<-w.compl #save a non-detrended version
year<-1961:2008

##get all the vectors
w.compl<-data.frame(t(apply(w.compl, 1,function(x) resid(loess(x~year))))) #detrend using loess fits. (doesn't matter if done local or global, but
residual<-(colSums(w.compl)) #the residual on global scale
actual.mean<-predict(loess(colSums(w.compl.nd)~year)) #the mean observed valuenon global scale
perc.dev<-residual/actual.mean #percent deviation
sd.resid<-sd(residual) #sd of the residuals
sd.diff<-residual/sd.resid #sd deviation

##call the values to output
sddiff.max<- min(sd.diff)
max.p.obs<-min(perc.dev)
max.year<-rownames(data.frame(perc.dev))[which.min(apply(data.frame(perc.dev),MARGIN=1,min))] #year of the maximum deficit

##join into a dataframe
 vars<-c(rep("sddiff.max",1), rep( "max.p.obs",1), rep( "max.year",1))
 data<-c(sddiff.max, max.p.obs, max.year)
 cropid<-c(rep(unlist(crop),3 ))
 all<-data.frame(vars,data, cropid) #crop specific
 all.crop.final<-rbind(all.crop.final, all)
}
```

```
all.crop.final

         vars                      data  cropid
1  sddiff.max         -1.79219129817371 soybean
2   max.p.obs         -0.14651191248488 soybean
3    max.year Soybean_1974_Area_ver8.nc soybean
4  sddiff.max          -2.9248021819215   maize
5   max.p.obs        -0.230305265375192   maize
6    max.year   Maize_1983_Area_ver8.nc   maize
7  sddiff.max            -3.563104982756    rice
8   max.p.obs        -0.083130154521226    rice
9    max.year    Rice_2002_Area_ver8.nc    rice
10 sddiff.max         -2.34540978346199   wheat
11  max.p.obs       -0.0793545059812062   wheat
12   max.year   Wheat_2003_Area_ver8.nc   wheat
```

##### 4.3.2 Scenario planning

Using the historical data, we constructed a worst-case scenario event under complete synchronization of production trends for each of the four crops, and compared expected losses under this setting to the losses expected under the observed trends. We set up our thought experiment to occur in the final year of the dataset, in 2008. To estimate the baseline losses, we used the number of standard deviations that the maximum losses fell over 1961-2008 (-1.8$\sigma$ for soybean, -2.9$\sigma$ for maize, -3.6$\sigma$ for rice and -2.3$\sigma$ for wheat), to gain the lower bounds of a 100% historical prediction interval for productivity of this period. To estimate the losses under the worst-case scenario (WCS), we estimated the inflation of the standard deviation in the data under synchrony using the variance-covariance matrix of production trends, i.e. $\sigma_G^2 = (\sum_{i=1}^{n} \sqrt{x_{i,i}})^2$, and multiplied this by these scaling factors for each crop to obtain a maximum negative deviation from the mean under synchrony.

As we have identified two scenarios ('Baseline' and 'Worst Case Scenario'), above, we run four mitigation scenarios under the 'worst case scenario', to see what influence practical actions could have on production deficits. We split these mitigation strategies into variance reducing strategies, and mean increasing strategies. Both of these strategies work to reduce the expected minima of global production, but they work in different ways.

Variance reducing strategies work by making the system more stable overall, whereas mean increasing strategies, simply work to reduce the size of the deficit when expressed as the percent deviation from mean annual production trends. Both types of strategies are grounded in realistic interventions. For example, variance reduction strategies, can be implemented by diversifying genotypes, by adapting climate smart cropping systems, and by using either ecological engineering or developing technological infrastructure to resist environmental stressors. Importantly we might focus on doing this over the whole world, or we might focus on doing this only in the worlds breadbaskets. We will assess the impact of each. Mean increasing techniques can be achieved through expansion of agricultural land, or more palatably with respect to biodiversity loss, through increasing yield ceilings and decreasing yield gaps. Like variance reducing interventions, mean increasing techniques are also largely spatially dependent. The scenarios we are analyzing are therefore:

1. The observed time series, with no manipulation
2. The observed time series, with the correlation between food producing cells set to 1, i.e. the 'worst case scenario' (WCS)
3. WCS+ increasing local stability with a 50% reduction in variation in every grid cell across the world.
4. WCS+ increasing stability of breadbaskets, by reducing the variance of grid cells above the 90th percentile of contributers to the global production by 50% percent
5. WCS+ closing production gaps by increasing production of the grid cells <50th percentile of producing regions by 100%.

6. WCS+ raising production ceilings by increasing production in grid cells in top 90th percentile of producers by 100%.

Note, that below, we simply use make use of the fact that the standard deviation of global crop production, $\sigma_G$ under complete synchrony, when all elements of $P = 1$, is simply, $(\sum_{i=1}^{n} \sqrt{x_{i,i}})$, which is equal to the baseline worse case, and otherwise $\sigma_G$ is equal to $\sqrt{\sum_{i=1}^{n} \sum_{j=1}^{n} x_{i,j}}$.

```
rm(list = ls())#clear space
```

```
crops<-list("maize", "rice","soybean", "wheat")
all.crop.scen.te2=data.frame()
for(i in 1:length(crops)) {
  crop<-crops[i] #assign crop
   path=(paste(crop,"production100km2.rds", sep = "")) #create path
   w <- readRDS(path)  #read in
   w<- subset(w,, -c(x, y)) #remove co-ordinates
w.compl<-w[rowSums(is.na(w))!=ncol(w),] #remove complete cases of NA
w.compl[is.na(w.compl)] <- 0 # replace NA with zeros
w.compl.nd<-w.compl #save a non-detrended version
year<-1961:2008
w.compl<-data.frame(t(apply(w.compl, 1,function(x) resid(loess(x~year)))))#detrend the data

#baseline and worst cases (WCS)
  residual<-(colSums(w.compl)) #the residual on global scale
  actual.mean<-predict(loess(colSums(w.compl.nd)~year)) #the mean observed value on global scale
  sd.resid<-sd(residual) #sd of the residuals
  sd.diff.min<-min(residual/sd.resid)#sd deviation
  perc.dev<-(sd.resid*sd.diff.min)/actual.mean #percent deviation under baseline scenario 1
  sdlocal.p1<-sum(apply(w.compl,1,sd, na.rm=T)) #the sd under complete synchrony
  sync.resid<-sd.diff.min*sdlocal.p1 #the residuals under synchrony
  perc.dev.sync<-sync.resid/actual.mean #get the deviations under synchrony, i.e. WCS

#WCS+ variance reduction in all locations, by 50%
  sdlocal.p1.half<-sum(sqrt((apply(w.compl,1,sd, na.rm=T)^2)/2))
  scen3.resid<-sd.diff.min*sdlocal.p1.half #the residuals under scenario 3
  perc.dev.scen3<-scen3.resid/actual.mean #deviations under scenario 3

#WCS + variance reduction  in BB (top 90th percentile of producers) by 50%
  local.sum<-as.vector(rowSums(w.compl.nd, na.rm = FALSE, dims = 1)) #sum of production
  vars<-as.vector(apply(w.compl, 1, var, na.rm=T)) #variance in production
  quantile90<-ifelse(local.sum>=quantile(local.sum, probs=0.9), 1, 0) #top producers
  vars3<-ifelse(quantile90==1, vars*0.5, vars) #halve the variance of top producters
  sdlocal.p1.bb<-sum(apply(w.compl,1,sd, na.rm=T))
  sd.local.p1.bbvar<-sum(sqrt(vars3)) #get the sd
  scen4.resid<-sd.diff.min*sd.local.p1.bbvar #the residuals under scenario 4
  perc.dev.scen4<-scen4.resid/actual.mean #deviations under scenario 4

#WCS+ increase in largest producers by 100%
  local.sum<-as.vector(rowSums(w.compl.nd, na.rm = FALSE, dims = 1))
  quantile90<-ifelse(local.sum>=quantile(local.sum, probs=0.9), 1.5, 1) #identify top producers
  prod.scen5<-colSums(w.compl.nd*quantile90)
  scen5.mean<-predict(loess(prod.scen5~year))
  perc.dev.scen5<-residual/scen5.mean #percent deviation under scenario 5

#WCS+ increase smallest producers by 100%
  local.sum<-as.vector(rowSums(w.compl.nd, na.rm = FALSE, dims = 1))
  quantile50<-ifelse(local.sum<=quantile(local.sum, probs=0.5), 1.5, 1)
  prod.scen6<-colSums(w.compl.nd*quantile50) #bottom producers
  scen6.mean<-predict(loess(prod.scen6~year))
  perc.dev.scen6<-residual/scen6.mean #percent deviation under scenario 6

#call the values to output
   max.p.obs<-perc.dev[48]
   max.p.sync<-perc.dev.sync[48]
   max.p.scen3<-perc.dev.scen3[48]
   max.p.scen4<-perc.dev.scen4[48]
   max.p.scen5<-perc.dev.scen5[48]
   max.p.scen6<-perc.dev.scen6[48]

#join into a dataframe
   vars<-c(rep("1.Baseline",1),
           rep( "2.Synchronisation",1),
           rep("3.Global variance reduction",1),
           rep( "4.Bread-basket variance reduction",1),
           rep( "5.Closing production gaps",1) ,
           rep( "6.Raising production ceilings",1))
data<-c(max.p.obs, max.p.sync, max.p.scen3, max.p.scen4,max.p.scen5,max.p.scen6)
  cropid<-c(rep(unlist(crop),6 ))
  all<-data.frame(vars,data, cropid) #crop specific
  all.crop.scen.te2<-rbind(all.crop.scen.te2, all)
}
```

```
all.crop.scen.te2

                                vars   data  cropid
1                         1.Baseline -0.121   maize
2                  2.Synchronisation -0.363   maize
3        3.Global variance reduction -0.256   maize
4  4.Bread-basket variance reduction -0.297   maize
5          5.Closing production gaps  0.038   maize
6      6.Raising production ceilings  0.051   maize
7                         1.Baseline -0.076    rice
8                  2.Synchronisation -0.253    rice
9        3.Global variance reduction -0.179    rice
10 4.Bread-basket variance reduction -0.216    rice
11         5.Closing production gaps  0.039    rice
12     6.Raising production ceilings  0.050    rice
13                        1.Baseline -0.043 soybean
14                 2.Synchronisation -0.177 soybean
15       3.Global variance reduction -0.125 soybean
16 4.Bread-basket variance reduction -0.150 soybean
17         5.Closing production gaps -0.029 soybean
18     6.Raising production ceilings -0.037 soybean
19                        1.Baseline -0.077   wheat
20                 2.Synchronisation -0.352   wheat
21       3.Global variance reduction -0.249   wheat
22 4.Bread-basket variance reduction -0.309   wheat
23         5.Closing production gaps  0.063   wheat
24     6.Raising production ceilings  0.080   wheat
```

##### 4.3.3 Scenario plots

Next we finally plot everything out for the scenario analysis.

```
library(ggplot2)
scenariote2<-ggplot(all.crop.scen.te2, aes(x = vars, y=data*100, fill=vars))+
 geom_bar(stat="identity",width = 0.5)+
  theme_bw()+
  facet_wrap(~cropid)+
scale_fill_manual(values = c("#377eb8",  "#4daf4a", "#e41a1c","#ff7f00", "gray", "black"))+
  ylab(expression(paste("Production loss (%)")))+
  xlab("")+
  geom_hline(yintercept = 0, size = 0.4)+
 theme(plot.background = element_blank(),panel.grid.major = element_blank(), panel.grid.minor = element_blank(),
         legend.key = element_blank(),axis.text.x = element_blank())+
  theme(strip.text.y = element_blank())+
theme(strip.background = element_blank())+
   theme(panel.border= element_blank())+
  guides(fill = guide_legend(title = "Scenario", title.position = "top"), nrow=10)+
scale_x_discrete(labels=c("1" = "", "2" = "",  "3" = "", "4" = "", "5" = "", "6" = ""))+
        theme(axis.line.y = element_line(color="black", size = 0.4))+
  theme(axis.ticks.x=element_blank())
scenariote2
```

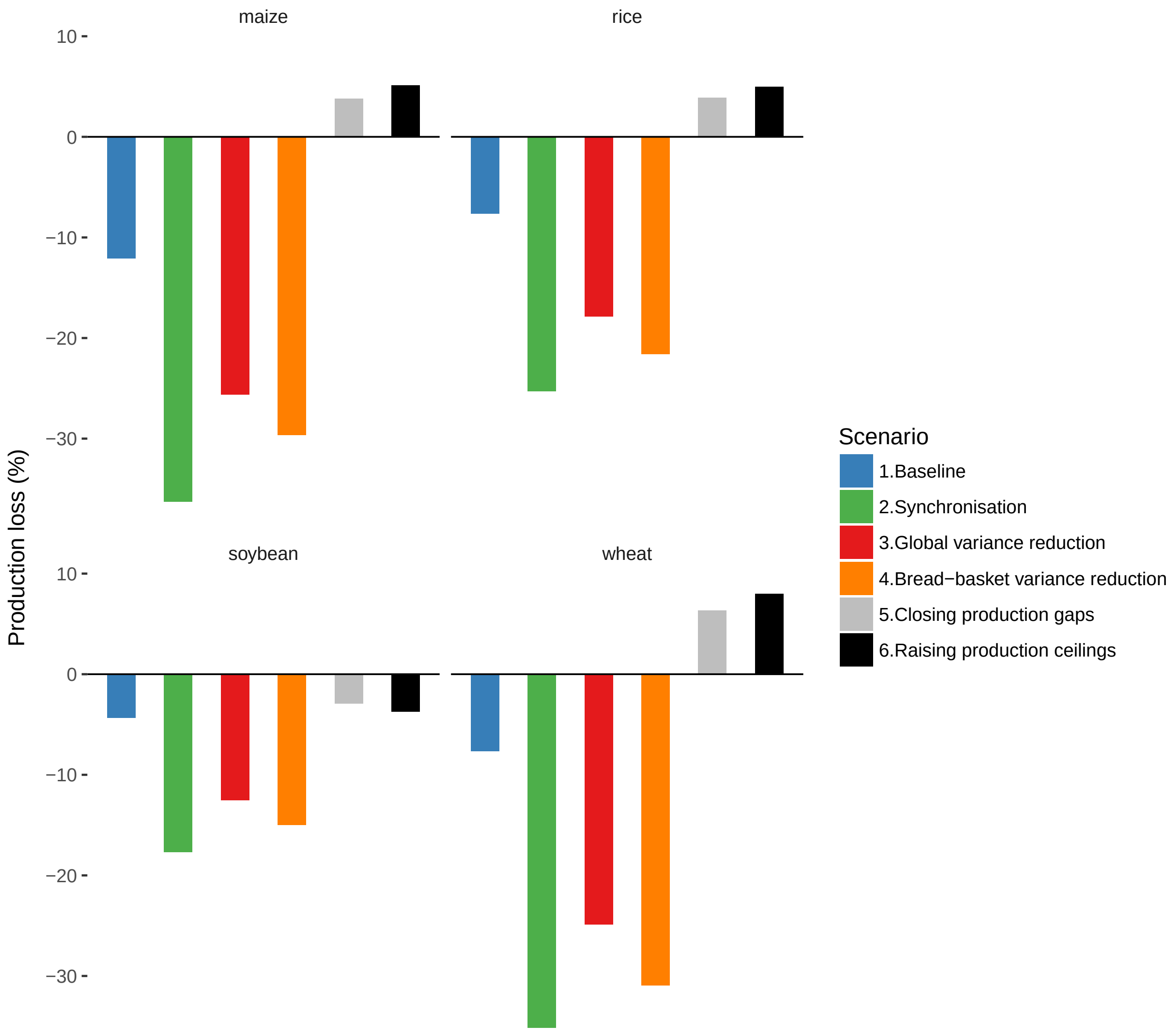

Figure 3: Strategies to cope with losses in a synchronised food system. Baseline= deviations from the mean for major crop commodities in 2008 are estimated using the lower bounds of the distribution of negative deviations from global trends over 1961-2008 (-1.8$\sigma$ for soybean, -2.9$\sigma$ for maize, -3.6$\sigma$ for rice and -2.3$\sigma$ for wheat). Syncronisation= worst case scenario (WCS), estimated by inflating baseline deviations by the variance increase due to complete syncrony in local production trends. Local variance reduction = WCS+ 50% reduction in variation in production for every grid cell across the world; Breadbasket variance reduction = WCS+ reducing the variance of grid cells in the 90-100th percentile of top producers by 50% percent; Closing production gaps= WCS+ increasing production of bottom 0-50th percentile of producers to 50%; Raising production ceilings= WCS+ increasing production in grid cells the 90-100th percentile of producers by 50%.

### Session information

```
sessionInfo()
```

```
R version 3.4.2 (2017-09-28)
Platform: x86_64-apple-darwin15.6.0 (64-bit)
Running under: macOS High Sierra 10.13.3

Matrix products: default
BLAS: /System/Library/Frameworks/Accelerate.framework/Versions/A/Frameworks/vecLib.framework/Versions/A/libBLAS.dylib
LAPACK: /System/Library/Frameworks/Accelerate.framework/Versions/A/Frameworks/vecLib.framework/Versions/A/libLAPACK.dylib

locale:
[1] en_CA.UTF-8/en_CA.UTF-8/en_CA.UTF-8/C/en_CA.UTF-8/en_CA.UTF-8

attached base packages:
[1] grid      stats     graphics  grDevices utils     datasets  methods
[8] base

other attached packages:
 [1] cowplot_0.9.2    gridExtra_2.3    ggplot2_2.2.1    rgdal_1.1-10
 [5] maptools_0.9-2   raster_2.6-7     sp_1.2-7         checkpoint_0.4.1
 [9] knitr_1.17       RevoUtils_10.0.6

loaded via a namespace (and not attached):
 [1] Rcpp_0.12.13     magrittr_1.5     munsell_0.4.3    colorspace_1.3-2
 [5] lattice_0.20-35  rlang_0.1.2      stringr_1.2.0    highr_0.6
 [9] plyr_1.8.4       tools_3.4.2      gtable_0.2.0     digest_0.6.12
[13] lazyeval_0.2.0   tibble_1.3.4     evaluate_0.10.1  labeling_0.3
[17] stringi_1.1.5    compiler_3.4.2   scales_0.5.0     foreign_0.8-69
```
